# Small Molecule Regulation of CLOCK:BMAL1 DNA Binding Activity

**DOI:** 10.64898/2026.04.13.718289

**Authors:** Diksha Sharma, Soumendu Boral, Ethen West, McClain Kressman, Irene Franco, Sarvind Tripathi, Hsiau-Wei Lee, Carlos A. Amezcua, Denize C. Favaro, Kevin H. Gardner, Carrie L. Partch

## Abstract

CLOCK:BMAL1 is a bHLH-PAS transcription factor complex that utilizes its bHLH (basic helix-loop-helix) domains to bind E-box motifs in DNA and tandem PAS (PER-ARNT-SIM) domains to heterodimerize and interact with regulatory proteins to generate circadian rhythms. PAS domains are evolutionarily conserved modules that frequently bind small molecule ligands within buried cavities to perform sensory and signal transduction functions. CLOCK and BMAL1 PAS domains have cavities that could be leveraged to regulate the transcription factor, and consequently, the circadian clock. Using NMR spectroscopy, we identified small molecules that bind within a cavity inside the PAS-A domain of CLOCK and its paralog NPAS2, which sits at an important flexible junction in the structured core of the heterodimer. We identified a gatekeeping mutant in the core of CLOCK PAS-A that significantly decreased ligand binding affinity. High-pressure NMR studies showed that ligand binding or the gatekeeping mutant significantly stabilized the domain. Finally, we showed that ligands induced dose-dependent displacement of CLOCK:BMAL1 from DNA *in vitro*. Together, these data demonstrate that small molecules can regulate DNA binding by the circadian transcription factor CLOCK:BMAL1 through occupancy of a PAS domain cavity.

## Introduction

Circadian rhythms are a product of several interlocking transcription/translation feedback loops that sustain ∼24-hour biological timekeeping [1, 2]. In most animals, circadian rhythms rely on the activity of the pioneer-like transcription factor CLOCK:BMAL1, which acts in the core feedback loop to drive the rhythmic expression of clock-controlled genes that regulate physiology and behavior [3–5]. Circadian dysregulation from genetic and environmental factors or aging contributes to cardiovascular and metabolic disorders, gut and immune dysfunction, and some cancers [6–8]. As the primary driver of animal circadian rhythms, CLOCK:BMAL1 is therefore an ideal target for pharmacological modulation to potentially combat these disorders by enhancing clock output [9, 10] or shutting down the clock in cancer cells that hijack circadian machinery for proliferation [11–13].

CLOCK:BMAL1 is a basic helix-loop-helix PER-ARNT-SIM (bHLH-PAS) transcription factor complex whose two subunits each contain three main elements: (1) an N-terminal bHLH domain that binds Enhancer-box (E-box) sites in DNA, (2) two tandem PAS domains (PAS-A and PAS-B) that form the structured core together with the bHLH domains, and (3) a disordered C-terminal region which recruits coactivators and repressors along that control CLOCK:BMAL1 activity [14–17]. PAS domains facilitate heterodimerization of BMAL1 with the ubiquitously expressed subunit CLOCK (Circadian locomotor output cycles kaput) or its paralog NPAS2 (Neuronal PAS domain-containing protein 2), as well as mediating direct contact with other core clock proteins [18, 19] and histones to form stable CLOCK:BMAL1:nucleosome complexes [20]. All of these PAS-based interactions are indispensable for CLOCK:BMAL1 function and circadian rhythms.

PAS domains are ubiquitous in all kingdoms of life, where they are found in many different types of sensory and signaling proteins. They typically have a conserved three-dimensional fold consisting of an antiparallel β-sheet flanked on one side by several *α*- helices, often generating a buried cavity between the helical and sheet layer that can bind chemically diverse small molecules [21]. PAS domains from bacteria, fungi, and plants frequently utilize this cavity to bind cofactors that sense environmental cues, such as light or oxygen, or to interact with small molecules for nutrient uptake [22–25]. By contrast, most PAS domains studied to date in animal proteins have an unoccupied or water-filled cavity and typically mediate protein-protein interactions [26, 27]. The aryl hydrocarbon receptor (AHR), a bHLH-PAS protein, is one apparent exception; its PAS-B domain binds a diverse array of xenobiotics, which trigger heterodimerization with the aryl hydrocarbon nuclear translocator (ARNT) and activation of a vast transcriptional program for xenobiotic detoxification [28, 29]. Across many species, PAS ligands trigger changes in conformation or dynamics to regulate protein function [23, 30, 31]. Based on this principle, orphan PAS domain-containing proteins have been targeted with exogenous small molecules, from PAS kinase (PASK) [32] to the bHLH-PAS subunits of hypoxia inducible factor (HIF), HIF2*α* and HIF1β/ARNT [33–36], and BMAL1 [37].

Here, we sought to identify small molecule ligands that could bind to the PAS-A domain of NPAS2 and its paralog CLOCK, regulating their biological DNA-binding activity in the process. We used a combination of NMR-based ligand screening and pressure perturbation to characterize ligand binding, both of which indicated that these ligands bind within the buried cavity of the PAS-A domain. We identified a gatekeeper mutation in CLOCK PAS-A that reduced ligand binding and stabilized the protein to pressure-induced unfolding*. In vitro* DNA binding assays demonstrated that ligand binding specifically disrupts E-box binding by the CLOCK:BMAL1 heterodimer. Although the top ligands here bind with modest micromolar affinity, this work provides a proof of principle that ligand binding within the buried cavity of CLOCK PAS-A can regulate DNA binding by CLOCK:BMAL1.

## Results

### NPAS2 PAS-A binds small-molecule ligands

CLOCK PAS-A is in a crucial position in the CLOCK:BMAL1 transcription factor, positioned immediately above a flexible pivot point between the bHLH and tandem PAS domains that allows the PAS domain core to undergo a rigid body rotation and stabilize nucleosome interactions that are necessary for circadian rhythms (Fig. 1A) [14, 20]. Restricting flexibility in PAS-mediated protein contacts can influence structural integrity and PAS function. For example, ligands that target HIF2*α* PAS-B induce rigidity across the domain and disrupt protein interactions with ARNT upon binding [35, 38]. Therefore, small molecules that bind within the PAS-A domain of CLOCK and its paralog NPAS2 have the potential to allosterically modulate the conformation and/or activity of the transcription factor to influence the circadian clock.

**Figure 1:**
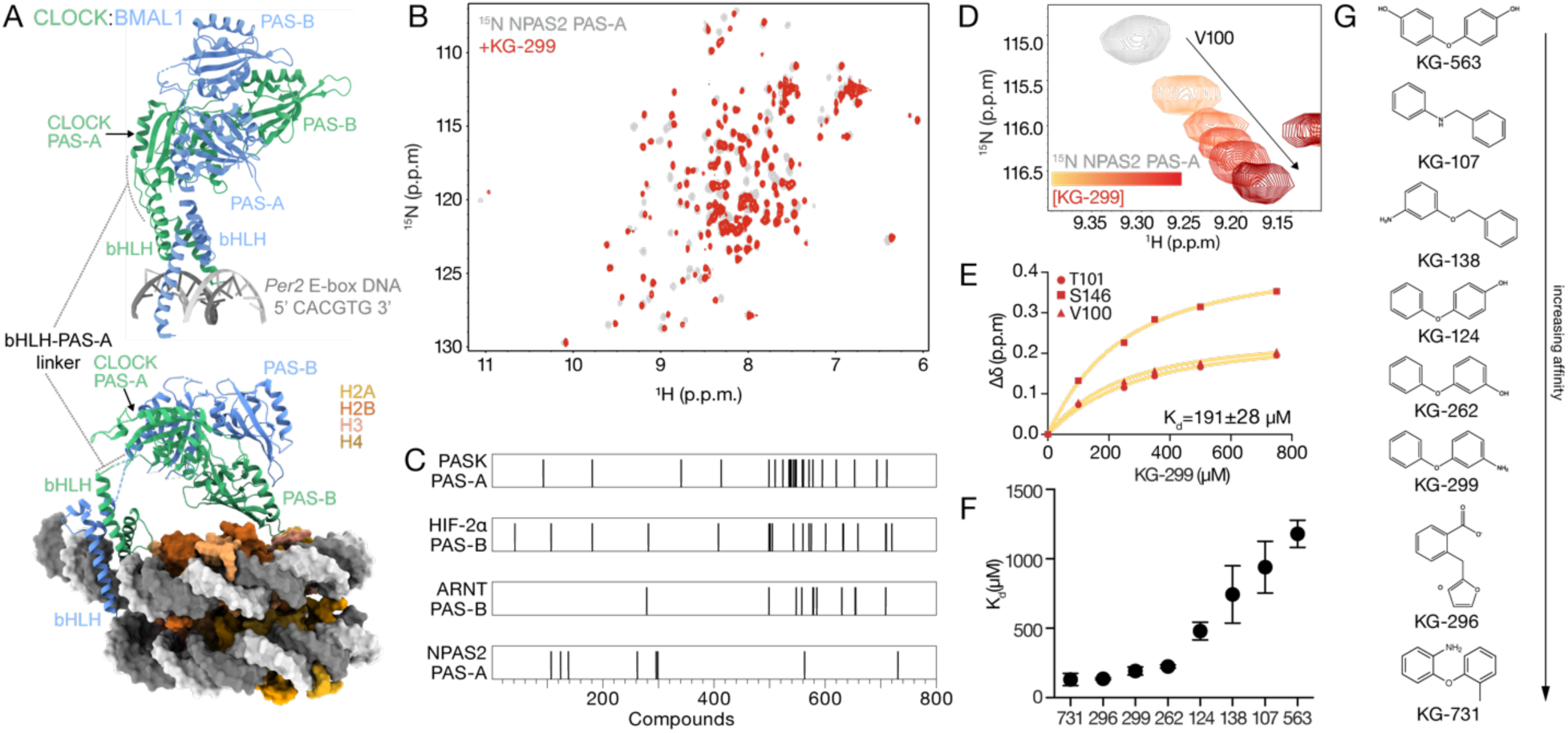
NMR ligand screening of NPAS2 PAS-A. **(A)** Overlay of crystal structures of CLOCK:BMAL1 bHLH-PAS-AB and bHLH bound to *Per2* E-box (PDB: 4F3L, 4H10) (top), CLOCK:BMAL1 bHLH-PAS-AB bound to nucleosome (PDB: 8OSK) (bottom). **(B)** ^15^N-^1^H HSQC spectra of ^15^N NPAS2 PAS-A (gray) with 500 μM KG-299 (dark red) **(C)** Hits from KG library for isolated PAS domains from different proteins. **(D)** Zoomed view of ^15^N-^1^H HSQC of NPAS2 PAS-A V100 (gray) showing dose-dependent response to KG-299. yellow to maroon (100-750 μM). **(E)** Fitting concentration-dependent CSPs on ^15^N labeled NPAS2 PAS-A to estimate K_d_ for KG-299. **(F)** Apparent K_d_ of 8 NPAS2 PAS-A hits from NMR screen ranked in order of affinity. Three residues were selected for each compound to fit K_d,_ and error bars represent fitted CSP errors. **(G)** Structures of 8 NPAS2 PAS-A hits from the NMR screen ranked in order of affinity.

We used an in-house library of 762 small molecules (average mass ∼200 Da) [32] that are well-suited to bind internal PAS cavities, which range from 100-600 Å^3^ in volume [21, 27]. We first aimed to identify small molecules that bind NPAS2 PAS-A due to the availability of backbone chemical shift assignments for the domain [39] that facilitated NMR screening and identification of ligand binding sites. Using ^15^N labeled NPAS2 PAS- A, we followed a standard NMR screening pipeline [32] and identified 8 ligands that induced chemical shift perturbations (CSPs) in the ^15^N-^1^H heteronuclear single quantum coherence (HSQC) spectrum of NPAS2 PAS-A, demonstrating direct binding to the protein (Fig. 1B and Fig. S1A). Prior NMR screens of this library against other isolated PAS domains yielded distinct hits from NPAS2 PAS-A (Fig. 1C) [32, 33, 36], demonstrating specificity at this early stage of screening. We observed that the NPAS2 PAS-A ligands all appear to target the same binding site, as they induced similar CSPs at identical residues, varying primarily in their apparent degree of binding (Fig. S1).

To rank the initial hits based on their affinity for NPAS2 PAS-A, we performed ligand titration experiments, collecting a series of ^15^N-^1^H HSQCs of ^15^N labeled NPAS2 PAS-A in the presence of increasing concentrations of each ligand. We observed fast exchange behavior in the titrations (Fig. 1D and Fig. S1B), exemplified by population-weighted average single peak moving linearly between apo and fully bound endpoints, consistent with affinities in the micromolar to millimolar range [40]. The CSPs for several residues from a representative ligand (KG-299) are shown in Fig. 1E (and others in Fig. S1C), which were fit to a one-site binding model to obtain an apparent dissociation constant (K_d_). The ranked hits (Fig. 1F) bound with the expected micromolar K_d_ affinities and share similarities in their structural scaffolds (Fig. 1G), likely explaining how they bind to the same region in NPAS2 PAS-A and cause similar chemical shift changes to largely the same subset of residues.

### Ligands bind within the buried cavity in CLOCK/NPAS2 PAS-A domain

Although NPAS2 can substitute for CLOCK in the suprachiasmatic nuclei (SCN) in the hypothalamus to sustain behavioral circadian rhythms [41, 42], CLOCK is more widely expressed in peripheral tissues and is therefore a more important target for circadian regulation. Human NPAS2 and CLOCK PAS-A domains are structurally conserved with an overall 64% sequence identity (Fig. S2A), but many of the residues lining the cavity are identical (Fig. S2B) [43]. Therefore, we hypothesized that ligands that bind NPAS2 PAS-A might also bind CLOCK PAS-A. We tested binding of three top NPAS2 PAS-A hits (KG-262, KG-296, KG-299) to ^15^N labeled CLOCK PAS-A by collecting ^15^N^-1^H HSQCs with ligand titrations. While KG-731 was another top hit like KG-296, we removed it from further consideration due to poor solubility. We found that KG-262, KG-299 and KG-296 bound CLOCK PAS-A with the same trend in affinity, with KG-296 having an apparent K_d_ of ∼120 μM (Fig. S2C). Hence, we selected KG-296 to biochemically characterize the effects of ligand binding on the NPAS2 and CLOCK PAS-A domains and in the CLOCK:BMAL1 heterodimer. We also performed a ligand-detected ¹H NMR experiment, WaterLOGSY, to independently confirm KG-296 binding to CLOCK and NPAS2 PAS-A. WaterLOGSY reports binding through the sign of the water-ligand NOE, which depends on molecular tumbling rate; a small, rapidly tumbling free ligand and a ligand bound to a slowly tumbling protein give NOEs of opposite sign, so a ^1^H resonance appears inverted relative to a non-binder [44, 45]. Fig. S2D shows the expected sign inversion for KG-296 in the presence of either CLOCK or NPAS2 PAS-A, confirming binding to both domains.

To determine the binding site of KG-296 on NPAS2 and CLOCK PAS-A, we quantified CSPs from ^15^N-^1^H HSQCs acquired during ligand titrations to both domains. Spectra of NPAS2 PAS-A (Fig. 2A) and CLOCK PAS-A (Fig. 2B) showed that both have several residues with large ligand-induced CSPs, which we quantified at saturating KG-296 concentrations at every assigned residue in both PAS-A domains. In both NPAS2 (Fig. 2C) and CLOCK (Fig. 2D), larger perturbations occurred primarily around two regions— the Aβ strand, AB-loop and F*α* helix on one side of the domain, and the Gβ and Hβ strands on the other. When we mapped these perturbations onto the structures of the PAS-A domains, we observed that KG-296 primarily influenced residues lining the CLOCK/NPAS2 PAS-A pocket (Fig. 2E). This region has high sequence identity in both PAS domains and across species, which explains their similar binding behavior with ligands (Fig. S2C,E). To test if KG-296 binds other PAS domains in the CLOCK:BMAL1 heterodimer, we collected ^15^N-^1^H HSQCs of ^15^N CLOCK PAS-B, BMAL1 PAS-A, and BMAL1 PAS-B with KG-296. We did not observe any ligand-dependent chemical shifts in these other PAS domains (Fig. S2D), consistent with high sequence divergence within the domains (Fig S2F). Altogether, our data indicated that KG-296 specifically binds within a buried cavity in CLOCK/NPAS2 PAS-A domain, which is the most conserved mechanism of ligand binding observed in other PAS-containing proteins with both endogenous [22–25] and exogenous ligands [32, 33, 35–37].

**Figure 2.**
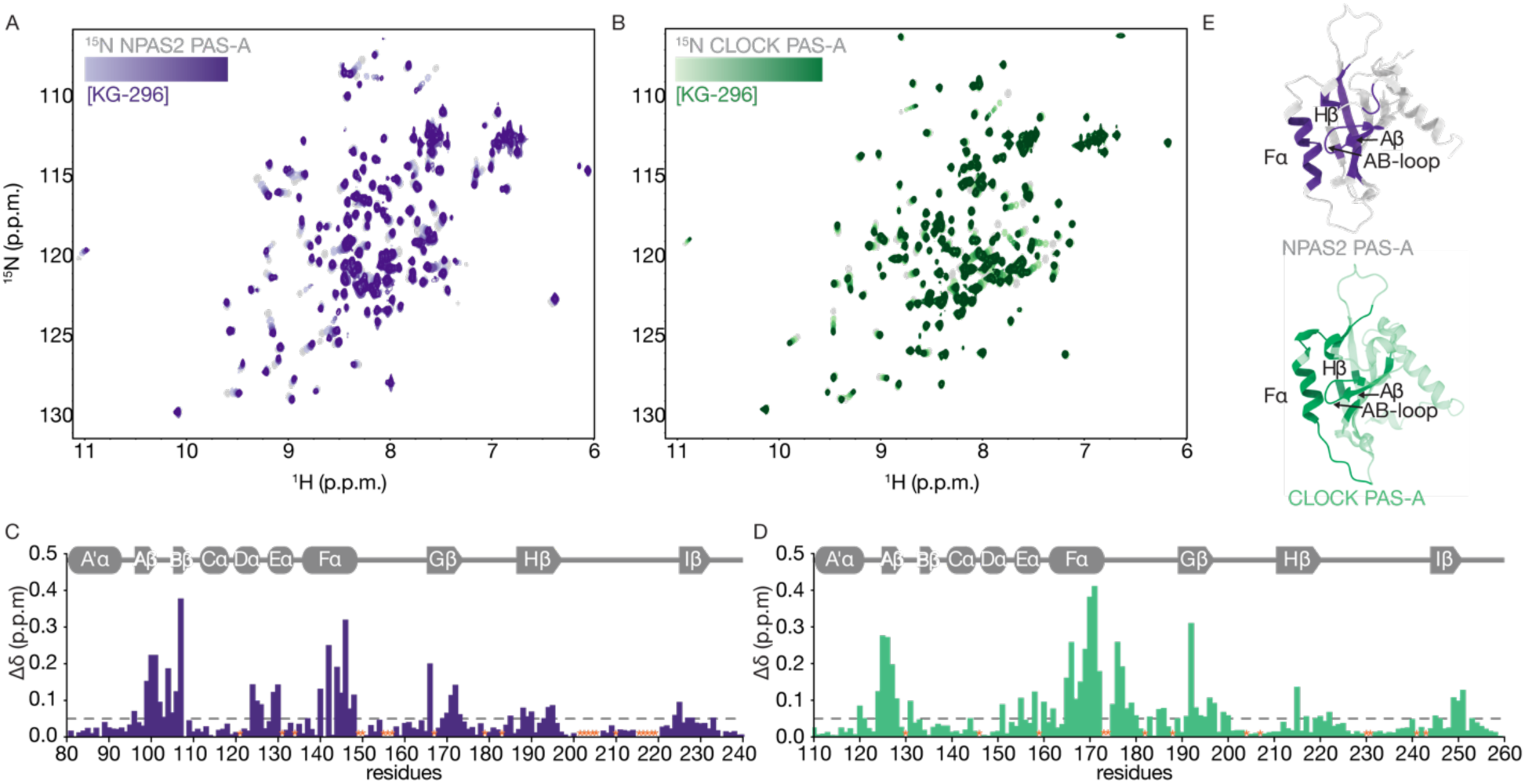
**KG-296 binds NPAS2 and CLOCK PAS-A within the cavity**. **(A)** ^15^N-^1^H HSQC spectra of ^15^N labeled NPAS2 PAS-A (gray) titrated with KG-296 (100-750 μM, light to dark purple). **(B)** ^15^N-^1^H HSQC spectra of ^15^N labeled CLOCK PAS-A (gray) titrated with KG-296 (100-750 μM, light to dark green). **(C)** Secondary structure of NPAS2 PAS- A (above) and CSP plot of ^15^N NPAS2 PAS-A with 1000 µM KG-296 (below). **(D)** Secondary structure of CLOCK PAS-A (above) and CSP plot of ^15^N CLOCK PAS-A with 1000 µM KG-296 (below). Dashed line in (C) and (D) indicates cutoff of 0.05 ppm. Orange asterisk, no data due to peak overlap or unassigned residues. **(E)** NPAS2 PAS-A (AlphaFold) with CSP > 0.05 ppm due to KG-296 (purple, top), CLOCK PAS-A (PDB 6QPJ) with CSP >0.05 ppm due to KG-296 (green, bottom).

### Gatekeeping mutant attenuates ligand binding

CLOCK PAS-A has an buried cavity that can be occupied by small-molecule ligands (Fig. S3A) [46]. To test if KG-296 binds inside this buried cavity of CLOCK PAS-A, we reasoned that introducing a larger side chain into the cavity might disrupt ligand binding. For example, a single point mutant inside the cavity of HIF2*α* PAS-B attenuated ligand affinity by 40-fold [47], and computational repacking of the core that shrunk the cavity size by 70% completely abrogated ligand binding [48]. To select mutations in CLOCK PAS-A, we used Pythia [49], which estimates protein stability changes (ΔΔG) caused by systematic mutation across the protein. We prioritized mutants based on three criteria: (1) the residue must be present in the binding site mapped by NMR (i.e., F*α* helix, AB-loop, or Hβ strand), (2) the mutant should have a bulkier side chain, and (3) the ΔΔG score should be less than 5 kcal/mol. We selected 8 mutants, out of which 4 (T127K, L170F, F193Y, C195F (Fig. S3B-C) yielded sufficient quantities of soluble protein for NMR analyses. All mutants had well-resolved NMR spectra comparable to native CLOCK PAS-A with modest mutation-associated CSPs (Fig. S3D). We collected ^15^N-^1^H HSQC spectra of each mutant with KG-296 and monitored ligand-dependent chemical shifts to assess binding. T127K, F193Y and C193F showed ligand-induced shifts similar to WT CLOCK PAS-A, while L170F had shifts that were decreased in magnitude compared to WT, suggesting weakened binding to KG-296 (Fig. S3E). We then titrated ^15^N CLOCK PAS-A L170F with KG-296 (Fig. 3A-B), revealing that residues near the F*α* helix and AB-loop exhibited the largest CSPs in the PAS-A domain, similar to WT. We calculated a ∼3.7-fold reduction in affinity (apparent K_d_ ∼ 444 μM, Fig. 3C), with diminished perturbations across the domain in the L170F mutant compared to WT CLOCK PAS-A (Fig. 3D).

**Figure 3.**
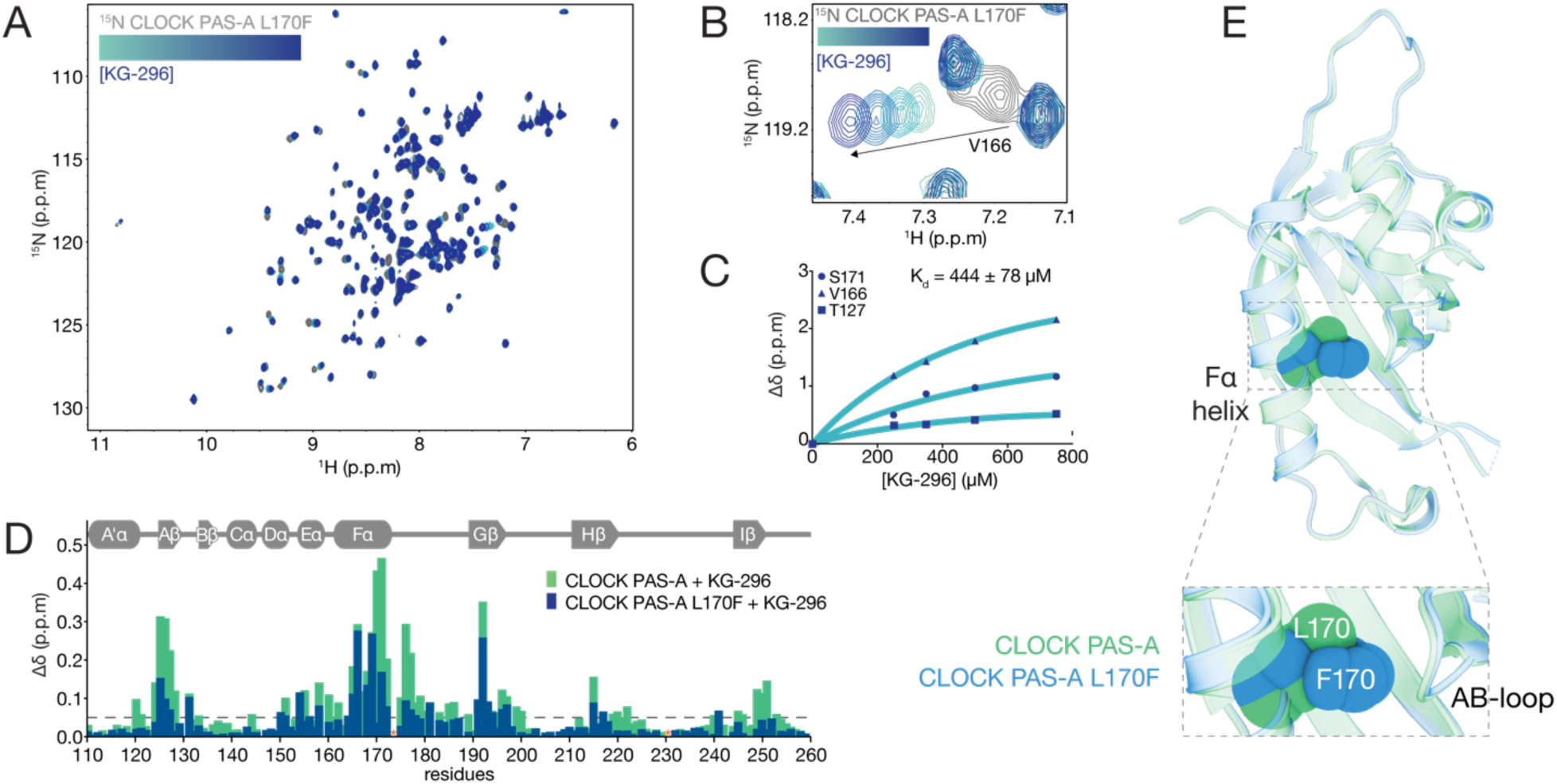
**Gatekeeping mutant attenuates KG-296 binding to CLOCK PAS-A**. **(A)** ^15^N- ^1^H HSQC spectra of ^15^N labeled CLOCK PAS-A L170F (gray) titrated with KG-296 (100- 750 μM, turquoise to dark blue). **(B)** Zoomed view of ^15^N-^1^H HSQC of V166 in CLOCK PAS-A L170F (gray) showing dose-dependent response of KG-296 (100-750 μM, turquoise to dark blue). **(C)** Fitting concentration-dependent CSPs on ^15^N labeled CLOCK PAS-A L170F to estimate apparent Kd for KG-296. **(D)** Secondary structure of CLOCK PAS-A WT/L170F (above) and overlay of CSP plot of ^15^N labeled CLOCK PAS-A WT (green) and L170F (dark blue) with 1000 µM KG-296. Dashed line, cutoff of 0.05 ppm. Orange asterisk, no data due to peak overlap or unassigned residues. **(E)** Structure overlay of CLOCK PAS-A WT (green, PDB 6QPJ) and CLOCK PAS-A L170F (dark blue, PDB 9PTN); F170 and L170 are shown in spheres.

To explore why the L170F mutant decreased KG-296 binding, we solved a crystal structure of the mutant (Table S1) and observed that introduction of the bulky phenylalanine side chain likely hinders access to the prospective binding region between the F*α* helix and AB-loop (Fig. 3E). Therefore, L170F acts as a gatekeeping mutant that obstructs KG-296 interaction with the internal cavity of the PAS-A domain. Despite numerous attempts to obtain a structure of KG-296 bound to wild-type CLOCK PAS-A, we could only partially resolve density for the ligand in the core of the protein, as validated by the composite omit map in electron density data. (Fig. S3F). We observed that the overall PAS-A fold with KG-296 was largely preserved with localized conformational shifts in the backbone and side chains of residues lining the F*α* and AB-loop region. We found that saturating amounts of ligand induced measurable backbone rearrangements in the Fα helix residues (L170, Y167), which shift modestly (0.6–2.0 Å), while the AB-loop (T127, D128, G129) undergoes a larger ∼2.1 Å displacement, suggesting the AB-loop repositions to accommodate the ligand into the PAS-A cavity (Fig. S3G).

### Ligand binding stabilizes CLOCK PAS-A under pressure perturbation

To further explore changes in CLOCK PAS-A upon KG-296 binding, we turned to high-pressure solution NMR spectroscopy. High-pressure perturbations to proteins are used to study folding-unfolding landscapes, explore low-lying excited states that are otherwise invisible at ambient pressure, and determine the effect of internal cavities on protein stability [50–55]. Higher hydrostatic pressure applied to proteins perturbs internal cavities and protein unfolding, which can be easily assessed by NMR [55, 56]. Studies have used high-pressure NMR to identify and study the stability effects of ligand binding on void-containing proteins, including PAS domains [54, 57]. Based on these principles, we were interested in investigating how the gatekeeping mutant and KG-296 binding affected the CLOCK PAS-A domain.

To assess the effects of pressure on the CLOCK PAS-A domain, we collected a series of ^15^N-^1^H HSQC spectra in a zirconium tube connected to an external pump allowing for rapid and reversible jumps in sample pressure. First, a baseline HSQC spectrum of ^15^N CLOCK PAS-A WT and the L170F mutant was collected at 20 bar, with and without saturating amounts of KG-296; then HSQC spectra were acquired after jumps to increasingly 250-2500 bar pressures, allowing us to track the effects of pressure on a residue-specific level across the protein (Fig. S4A). After each pressure jump, the pressure was lowered to 20 bar, allowing us to acquire spectra to assess the reversibility of pressure-induced conformational changes (Fig. 4A). Individual HSQC spectra at each increased pressure point and interleaved 20 bar basal pressure were acquired for 1 hour, resulting in a full pressure series spanning about 21 hours. In Fig. 4A, the x-axis reflects cumulative acquisition time, with each interleaved 20-bar point reporting the extent of peak intensity recovery after the preceding pressure jump. This analysis revealed a significant loss of the WT peak intensities from exposure to 2250 bar compared to the gatekeeping mutant L170F (Fig. 4B). Ligand binding also significantly stabilized both the WT and L170F mutant relative to their apo states, although ligand binding had a larger stabilizing effect on the WT protein (Fig. 4B). The irreversible effects of pressure on the stability of the WT protein are easily seen in the ^15^N-^1^H HSQC spectra; in contrast to the well dispersed peaks of the native fold found in the initial baseline spectrum (Fig. 4C, gray), the baseline spectrum taken after the jump to 2250 bar (Fig. 4C, black) shows a collapse of amide ^1^H chemical shift dispersion, consistent with an unfolded state. The addition of ligand to the CLOCK PAS-A domain or partial filling of the cavity with the L170F mutation reduced or eliminated the appearance of the unfolded state (Fig. 4C, green or yellow, respectively). For clarity, horizontal slices taken at ∼118 ppm ^15^N chemical shift show the loss of peak intensities at different pressure states (Fig. S4B). Taken together, these data demonstrate that KG-296 significantly stabilizes the CLOCK PAS-A domain at high pressure, confirming occupancy of the internal cavity.

**Figure 4.**
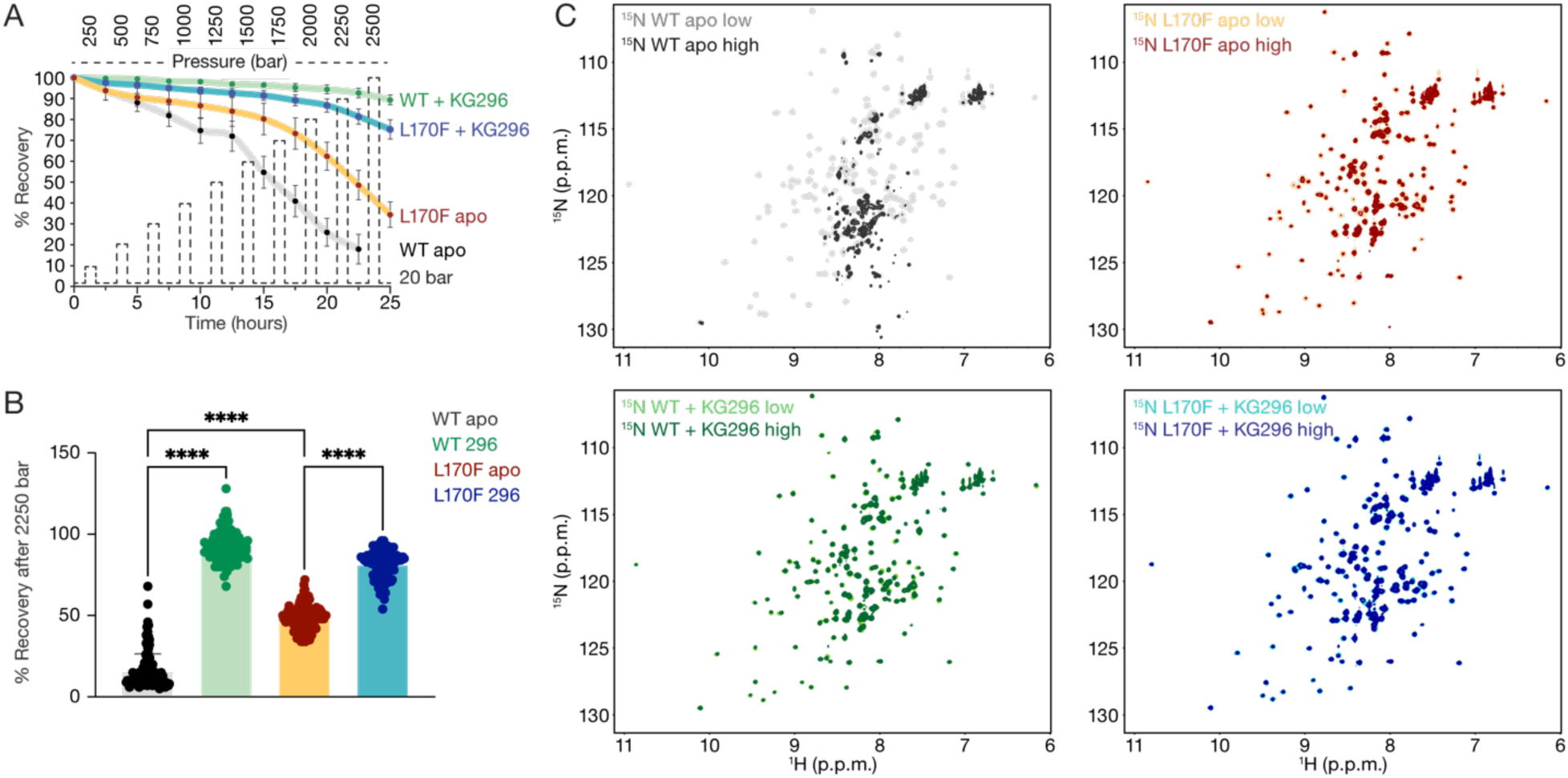
KG-296 stabilizes CLOCK PAS-A under high-pressure perturbation. **(A)** Graphical representation of average of peak intensities at 20 bar after individual pressure titration points. WT apo (gray), WT + 1200 µM KG-296 (green), L170F apo (mustard), L170F + 1200 µM KG-296 (turquoise). **(B)** Bar graph showing average peak intensities at 20 bar after the 2250 bar pressure point. Significance was calculated using a one-way ANOVA statistical test; ***, p<0.001; ****, p<0.0001. (A,B) Number of peaks used for mean ± s.d. calculation for WT apo/WT+KG-296 = 130; L170F apo = 129; L170F+KG- 296 = 124. **(C)** Overlay of ^15^N-^1^H HSQC spectra at 20 bar beginning of pressure titration point 250 bar (low) and at 20 bar after end of pressure titration point 2250 bar (high). WT apo (low, gray; high, black—top left), CLOCK PAS-A + 1200 µM KG-296 (low, light green; high, dark green—bottom left), L170F apo (low, yellow; high, maroon—top right), L170F + 1200 µM KG-296 (low, turquoise; high, dark blue—bottom right).

### Ligand binding inhibits DNA binding to CLOCK:BMAL1

Ligands that target PAS domains in bHLH-PAS transcription factors have diverse effects on function; they disrupt heterodimerization and DNA binding [36, 38], inhibit interactions with coactivators [33, 58], or modulate activity in a way that is apparently independent of DNA binding [37]. While the modest affinity of KG-296 for CLOCK PAS-A precluded its use in cellular assays, we assessed the effect of ligand binding on DNA binding of CLOCK:BMAL1 *in vitro*. First, we performed electrophoretic mobility shift assays (EMSAs) using bHLH and tandem PAS domains (bHLH-AB) fragments of CLOCK:BMAL1 WT and L170F to investigate binding to a Cy5-labeled 19-base pair double-stranded DNA probe from the mouse *Per1* promoter with the E-box sequence CACGTG. Both WT and L170F bound the DNA similarly (Fig. 5A, B). We also determined that WT and L170F CLOCK activate the transcription of a mouse *Per1*-luciferase reporter and express similarly when co-expressed with BMAL1 in HEK293T cells (Fig. 5C, D). Therefore, the L170F mutation does not appear to influence CLOCK:BMAL1 DNA binding *in vitro* or activity in cells.

**Figure 5.**
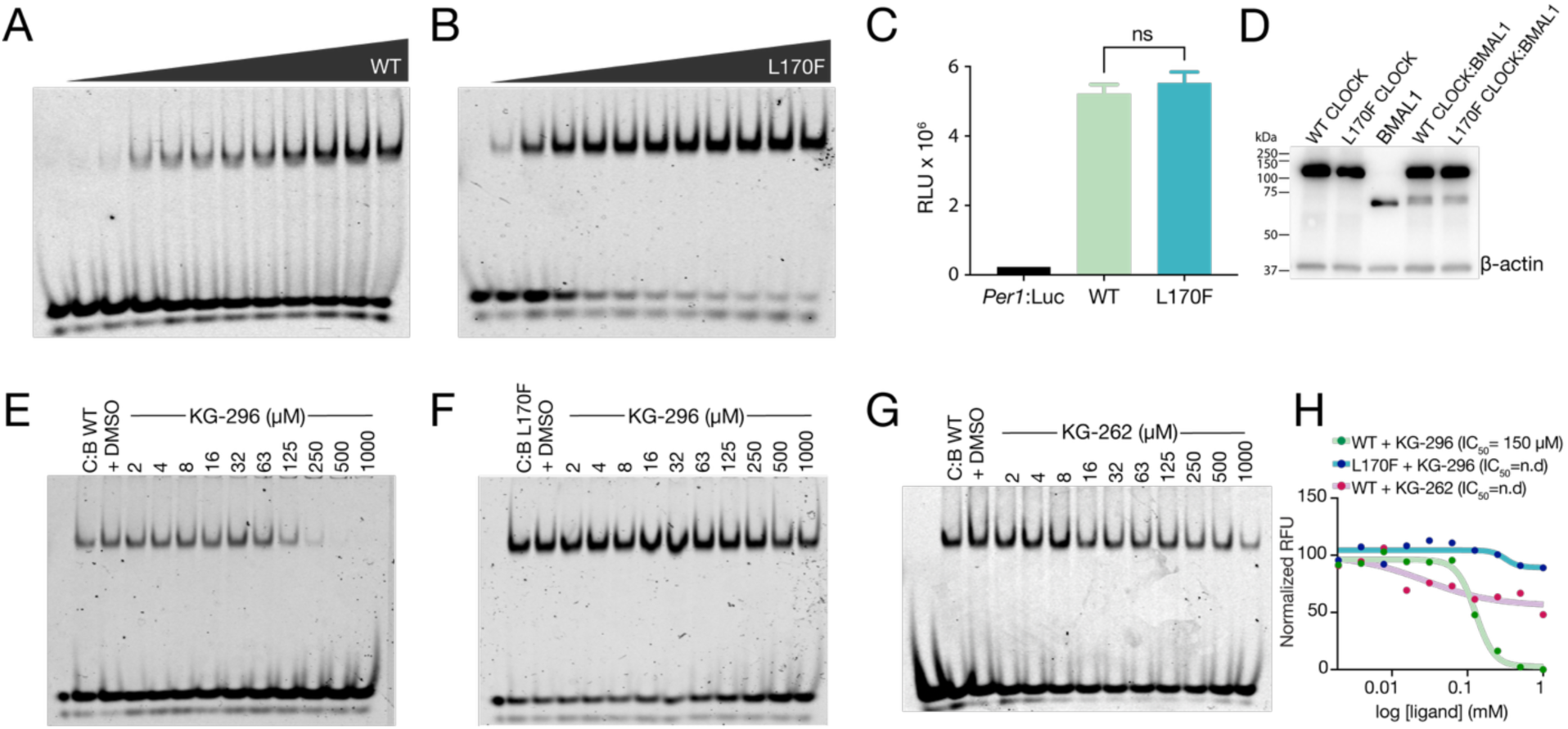
KG-296 inhibits DNA binding to CLOCK:BMAL1. (A-B) EMSA titration assay between CLOCK:BMAL1 (C:B) **(A)** WT (0-1000 nM) or **(B)** L170F (0-1000 nM) and E-box (4 nM). **(C)** *Per1*:luciferase reporter assay in HEK293T cells transfected with CLOCK:BMAL1 WT (light green) or CLOCK:BMAL1 L170F (turquoise). n = 8, mean ± s.d. Significance was calculated using one-way ANOVA statistical test, n.s., not significant. **(D)** Western blot of CLOCK:BMAL1 WT and L170F expressed in HEK293T cells with β- actin used as loading control. n = 2. **(E)** EMSA competition assay between CLOCK:BMAL1 WT and 0-2000 µM KG-296, **(F)** CLOCK:BMAL1 L170F and 0-1000 µM KG-296, **(G)** CLOCK:BMAL1 WT and 0-1000 µM KG-262. Three independent experiments of each of **(E-G)** were performed, and one representative gel is shown of each. **(H)** Normalized RFU from densitometry analysis of (E-G) plotted against ligand concentration. WT + KG-296, green; L170F + KG-296, blue; WT + KG-262, pink.

We then examined the effect of KG-296 on DNA binding by CLOCK:BMAL1 by forming a complex of the heterodimer with E-box DNA and titrating in KG-296 or DMSO alone. We observed a dose-dependent decrease in the DNA-bound complex with KG-296 that is consistent with the calculated K_d_ of KG-296 binding to CLOCK PAS-A, demonstrating that KG-296 disrupts CLOCK:BMAL1 binding to DNA with an IC50 of 150 μM (Fig. 5E,H). We performed the same KG-296 titration with the L170F mutant and observed that the protein-DNA complex largely remained intact, even at the highest ligand concentrations we tested (1 mM KG-296, Fig. 5F), consistent with the reduced affinity of the L170F variant for KG-296. We then performed an EMSA on WT CLOCK:BMAL1 bHLH-AB, titrating up to 1 mM of the low-affinity ligand KG-262 (K_d_ = 606 ± 90 μM) and, as expected, observed lower efficacy in inhibiting CLOCK:BMAL1 binding to DNA compared to KG- 296 (Fig. 5G). Altogether, these data demonstrate that binding of KG-296 into the internal buried cavity in CLOCK PAS-A specifically disrupts DNA binding by CLOCK:BMAL1.

## Discussion

PAS domains form the core structured region in the transcription factor CLOCK:BMAL1. They are essential for several inter- and intramolecular protein-protein interactions [59]: their dimerization is required for DNA binding [14, 15], they directly anchor to histones for stable nucleosome interactions [20], they bind to repressor proteins to maintain circadian periodicity [19], and undergo post-translational modifications that may influence the timing of the clock [60]. Many PAS domains have a buried cavity which binds chemically diverse small molecule ligands to regulate PAS domain activity, as seen in a range of prokaryotic and eukaryotic signaling proteins [28, 36, 61]. Therefore, leveraging the binding cavity in CLOCK:BMAL1 PAS domains for pharmacological regulation of the transcription factor could be a valuable tool to probe its control of the circadian clock and lay the foundation for therapeutic approaches to tackle circadian dysfunction in humans.

Here, we used NMR spectroscopy to identify small molecule ligands that bind to the PAS- A domain of NPAS2 and its paralog CLOCK. We found that KG-296, with a size of 202 Da, binds with moderate affinity within the conserved buried cavity in CLOCK/NPAS2 PAS-A and disrupts DNA binding by CLOCK:BMAL1 *in vitro*. We identified a gatekeeping mutant in CLOCK PAS-A, L170F, that reduces access of KG-296 to the cavity, attenuating binding. We used high-pressure NMR to probe the thermodynamic consequences of ligand binding to PAS domains and observed that KG-296 binding renders CLOCK PAS-A more stable and resistant to pressure-induced unfolding. Altogether, this work establishes that it is possible to directly regulate DNA binding by the core mammalian circadian transcription factor, CLOCK:BMAL1 by targeting a PAS domain cavity.

While other protein/ligand screening methods offer higher throughput, protein-detect NMR is ideal for studying weak interactions at a residue-specific level, allowing us to simultaneously detect potential ligands and identify the binding site. Our findings from the NPAS2/CLOCK PAS-A screen showed that the hits shared chemical similarities and bound to the same region in the protein with varying affinity. These differences will help rationalize future SAR (Structure Activity Relationship) efforts for hit optimization.

Furthermore, the specificity of KG-296 for the NPAS2/CLOCK PAS-A over all other PAS domains in CLOCK:BMAL1 transcription factor is a promising start toward drug development. This specificity is attributed to the particularly high diversity in amino acid sequence among PAS domains, an evolutionary mechanism that allows PAS domains to bind a diverse range of cofactors and small molecules despite maintaining a conserved topology [21, 61].

Our NMR data revealed that KG-296 induces maximum perturbations around the F*α* helix and AB-loop, along with some of the interior of the β-sheet, suggesting that KG-296 binds to the CLOCK PAS-A internal cavity. To validate the binding site, we introduced a L170F mutant on the F*α* helix right across from the AB-loop and found that it reduces KG-296 binding to CLOCK PAS-A, likely due to the bulkier side chain hindering the binding site. MD simulations have suggested two primary pathways for ligand entry and exit into the PAS-B domain of HIF2*α* transcription factor [62, 63]. Both pathways involve the F*α* helix (either between F*α*-Gβ or F*α*-AB-loop) in HIF2*α* PAS-B, which undergoes changes that likely allow the ligand in and out of the cavity. Therefore, it is possible that L170F acts by limiting access to the ligand entry site through the helical face of the PAS domain.

We were unsuccessful in obtaining a crystal structure of KG-296 bound within CLOCK PAS-A. We attribute this to the moderate affinity of KG-296, limiting the occupancy of this ligand within the protein, preventing us from obtaining the strong (2mF_o_−DF_c_) density needed to confidently model its location in the structure. However, we observed backbone and sidechain displacements of CLOCK PAS-A residues by up to 2.1 Å around the F*α* helix and the AB-loop in crystals soaked with saturating amounts of KG-296, corroborating our NMR CSP mapping data. Our high-pressure NMR data also support this model, as KG-296 stabilizes CLOCK PAS-A from pressure-induced unfolding. Such increased stability of protein molecules under high hydrostatic pressure is observed in structurally compact proteins with lower void volumes [50, 56]. Therefore, the increased stability of KG-296-bound CLOCK PAS-A suggests that it has a reduced void volume compared to the apo state as would be expected from interior ligand binding. Taken together, our data provide structural evidence of KG-296 binding into the CLOCK PAS-A cavity, supporting a conserved mechanism of PAS-ligand interaction.

We tested for a functional effect of KG-296 on CLOCK PAS-A in a more native context, using the larger, multidomain structured core of the CLOCK:BMAL1 heterodimer that binds to E-box DNA [14]. We found that KG-296 causes dissociation of a complex containing the bHLH and tandem PAS domains of CLOCK:BMAL1 with DNA, indicating that KG-296 has the potential to regulate CLOCK:BMAL1 in a different manner from a recently described BMAL1 PAS-B ligand [37]. We confirmed that this effect is due to binding to the CLOCK PAS-A domain, because it is reduced with the L170F gatekeeper mutant or with a lower-affinity ligand. Although both WT and L170F perturb the same region of the PAS-A domain, restricting access to the cavity is not equivalent to occupying it, which explains why L170F leaves DNA binding intact while KG-296 disrupts it. Consistent with this, KG-296 fails to displace the L170F complex from DNA (Fig. 5F), indicating that the inhibitory effect requires access to the PAS-A cavity. Unfortunately, the modest affinity of KG-296 for CLOCK PAS-A did not permit us to study the consequences of disrupting the interaction with DNA in cells. Discovery of a higher-affinity ligand with more amenable pharmacokinetic properties will undoubtedly give us better insights into the potential inhibitory effect of CLOCK PAS-A ligands on CLOCK:BMAL1 function.

Due to the vast influence of circadian rhythms on physiology and health, clock-targeting molecules are being sought to pharmacologically modulate the molecular clock [64]. CLOCK:BMAL1 presents itself as a crucial target due to its activator function in the core TTFL that drives animal circadian rhythms. However, regulating CLOCK:BMAL1 activity could have either enhancing or attenuating effects on the robustness, or amplitude, of circadian rhythms. CLK8, a compound that binds to bHLH region of CLOCK, disrupts BMAL1 binding, yet somehow enhances circadian amplitude in cellular assays [10]. By contrast, binding of the compound CCM to the internal cavity of BMAL1 PAS-B somehow decreases circadian amplitude without affecting DNA binding or CLOCK:BMAL1 heterodimerization [37]. Our study shows that KG-296 likely has an inhibitory mode of action on CLOCK:BMAL1 by disrupting its binding to DNA. This could be beneficial in glioblastoma cancer stem cells that rely on CLOCK:BMAL1 activity to proliferate, where indirect repression of CLOCK:BMAL1 retards cancer growth [11–13]. Generating analogs of KG-296 or other hits with enhanced affinity will enable a better understanding of the structural, biochemical, and cellular mechanisms of small molecule binding to clock protein PAS domains.

## Acknowledgments

Support for this work was provided by US National Institutes of Health grants R35 GM141849 (C.L.P.) and R35 GM156296(K.H.G.), and the Howard Hughes Medical Institute (C.L.P.). D.S. was supported by a predoctoral fellowship from the California Institute for Regenerative Medicine (CIRM) under award number EDUC4-12759 and the Institute for the Biology of Stem Cells (IBSC) at UC Santa Cruz. We thank James Partch for his contribution to figure designs and artwork.

## Author contributions

D.S., C.A., K.H.G., C.L.P. conceived of the study; D.S., E.W., S.T., H.-W.L., S.B., M.K., I.F., S.T., C.A., and D.C.F. performed experiments and analyzed results; K.H.G., C.L.P. oversaw the work and secured funding; D.S. and C.L.P. wrote the manuscript with edits from the other authors.

## Materials availability

Materials are available upon request from Carrie Partch or Kevin H. Gardner with a materials transfer agreement from their respective institutions.

## Data availability

The coordinates for CLOCK PAS-A L170F are deposited in the RCSB Protein Data Bank (PDB: 9PTN). NMR chemical shift assignments for CLOCK PAS-A wild-type and L170F are deposited in the Biological Magnetic Resonance Data Bank (BMRB: 53362 and 53377, respectively).

## Materials and methods

### Protein expression and purification

#### NPAS2 and CLOCK PAS-A

Mouse NPAS2 PAS-A (UniprotKB P97460) (residues 78-240) and mouse CLOCK PAS- A (UniProtKB O08785) (residues 93-261) were cloned into a bacterial expression plasmid based on the pET22b vector backbone from the parallel vector series and expressed using *Escherichia coli* Rosetta2 (DE3) cells [65]. Gatekeeping point mutations in CLOCK PAS-A were generated using site-directed mutagenesis and verified by sequencing. Constructs have an N-terminal TEV-cleavable His_6_ or a His_6_-NusA extra-linker (HNXL) solubilizing tag and ampicillin resistance. Rosetta (DE3) *E. coli* harboring plasmids were grown to an OD600 of ∼0.6-0.9 at 37°C in the presence of ampicillin (100 µg/mL) and chloramphenicol (35 µg/mL). Protein expression was induced with 0.5 mM isopropyl β-D- 1-thiogalactopyranoside (IPTG) and allowed to proceed for 16-18 hours at 18°C in either Luria Broth (LB) or M9 minimal medium containing 1 g/L ^15^NH_4_Cl with or without 3 g/L ^13^C D-glucose to generate uniformly ^15^N-labeled or ^13^C,^15^N-labeled proteins for NMR spectroscopy. Cells were lysed with an Emulsiflex C-3 cell disruptor (Avestin) in Buffer A containing 50 mM Tris pH 7.5, 500 mM NaCl, 5 mM 2-mercaptoethanol, 5% glycerol and 20 mM imidazole. *E. coli* cell lysate was centrifuged at 19,000 rpm for 45 minutes to separate the soluble fraction from cell debris. The soluble fraction of *E. coli* lysate was passed over Ni-NTA resin (QIAGEN), washed thoroughly, and eluted using 250 mM imidazole. Fractions of interest were buffer exchanged into TEV lysis buffer using a HiPrep 26/10 desalting column (Cytiva). Proteolysis was performed with His_6_-tagged TEV protease overnight at 4°C and the cleaved protein was retained from the flow-through of a Ni-NTA column. The protein was further purified on a preparative grade Superdex 75 16/600 size-exclusion column (Cytiva) pre-equilibrated with NMR buffer (20 mM Tris-HCl pH 7.0, 50 mM NaCl, 2 mM TCEP). Purified protein was concentrated to the desired concentration using Amicon® Ultra centrifugal filter with a 10 kDa molecular weight cutoff (MWCO) (Merck Millipore).

Human CLOCK PAS-A (UniProtKB O15516 residues 106-265) was synthesized by Twist Bioscience for crystallography using the exact sequence from PDB 6QPJ. The DNA fragment was cloned into a bacterial expression plasmid based on the pET-29b(+) vector backbone from the parallel vector series with an N-terminal Ulp1-cleavable His_6_-GST- SUMO solubilizing tag and kanamycin resistance. The L170F mutation was generated using site-directed mutagenesis and verified by sequencing. Rosetta (DE3) *E. coli* harboring CLOCK PAS-A plasmids were grown to an OD600 of ∼0.6-0.9 at 37°C in the presence of kanamycin (100 µg/mL) and chloramphenicol (35 µg/mL). Protein expression was induced with 0.5 mM IPTG and allowed to proceed for 16-18 hours at 18°C in LB. Cells were lysed with an Emulsiflex C-3 cell disruptor (Avestin) in Buffer A containing 50 mM Tris pH 7.5, 500 mM NaCl, 5 mM 2-mercaptoethanol, 5% glycerol and 20 mM imidazole. *E. coli* cell lysate was centrifuged at 19,000 rpm for 45 minutes to separate the soluble fraction from cell debris. The soluble fraction of *E. coli* lysates was passed over Ni-NTA resin (QIAGEN), washed thoroughly, and eluted using 250 mM imidazole.

Fractions of interest were pooled, and proteolysis was performed with His_6_-tagged Ulp1 protease for 1 hour at 4°C. The cleaved protein was retained from the flow-through of a Ni-NTA column. The protein was further purified on a preparative grade Superdex 75 16/600 size-exclusion column (Cytiva) pre-equilibrated with crystallization buffer (20 mM Tris-HCl, 150 mM NaCl pH 7.5). Purified protein was concentrated to the desired concentration using Amicon® Ultra centrifugal filter with a 10 kDa MWCO (Merck Millipore).

#### CLOCK:BMAL1 bHLH-PAS-AB

Mouse CLOCK (UniProtKB O08785) bHLH PAS-AB (residues 26–395) and mouse BMAL1 (UniProtKB Q9WTL8-4) bHLH PAS-AB (residues 62–441) were cloned into separate pFastBac vectors as described previously [14]. The L170F mutation was generated in CLOCK using site-directed mutagenesis and verified by sequencing. 1–2 L of CLOCK-BMAL1 bHLH-PAS-AB-expressing insect cells (*Spodoptera frugiperda*) were pelleted and resuspended in His Buffer A (20 mM sodium phosphate buffer pH 8, 200 mM NaCl, 15 mM imidazole, 10% glycerol, 0.1% Triton X-100 and 5 mM 2-mercaptoethanol). Cells were lysed by cell disruption and subsequent sonication for 3 min on ice (15 sec on, 30 sec off). Lysate was clarified by centrifugation at 19,000 rpm at 4°C for 45 min. Ni-NTA affinity purification was performed on a 5 ml HisTrap FF affinity column (Cytiva). After 15- column washes in His Buffer A, the column was further washed with 6.5% His Buffer B (20 mM sodium phosphate buffer pH 7.5, 200 mM NaCl, 300 mM imidazole, 10% glycerol and 5 mM 2-mercaptoethanol) for 3 column volumes (CV) before being eluted in a 10 CV gradient to 100% His Buffer B. Fractions of interest were buffer exchanged into Heparin Buffer A (20 mM sodium phosphate buffer pH 7.5, 50 mM NaCl, 2 mM DTT and 10% glycerol) using a HiPrep 26/10 desalting column (Cytiva) and cleaved with His_6_-TEV at 4 °C overnight. The cleaved protein was loaded onto a HiTrap Heparin HP affinity column (Cytiva). After washing with 5 CV of the above buffer, the column was washed with a further 3 CV of 25% Heparin Buffer B (20 mM sodium phosphate buffer pH 7.5, 2 M NaCl, 2 mM DTT and 10% glycerol) before eluting with Buffer B over an 8 CV gradient. The protein was further purified on a preparative grade Superdex 200 16/600 size-exclusion column (Cytiva) pre-equilibrated with storing buffer (20 mM HEPES pH 7.5, 125 mM NaCl, 5% glycerol and 2 mM TCEP). Purified protein was concentrated to the desired concentration using Amicon® Ultra centrifugal filter with a 30 kDa MWCO (Merck Millipore).

### Selection of gatekeeping mutants in human CLOCK PAS-A

The structure of human CLOCK PAS-A (PDB:6QPJ) was uploaded on the Pythia web server (https://pythia.wulab.xyz/) [49]. The program substitutes every residue in the protein one at a time to the 20 common amino acids, predicts the mutant-driven stability, and calculates free energy changes (ΔΔG, difference in ΔG between WT and mutant. ΔΔG for mutants chosen for the study are C195F:0.44; F193Y:2.43; L170F:4.34; T127K:0.49 (all units in kcal/mol).

### NMR Spectroscopy

#### Small molecule screening

All NMR screening data were recorded at 25°C with Varian Inova 500 or 600 MHz spectrometers, using the pipeline described in [32]. Briefly, a cocktail of five compounds dissolved in d_6_-DMSO was added to ^15^N labeled NPAS2 PAS-A (250 μM), resulting in the final concentration of each ligand at 1 mM. Potential hits were identified by comparing the chemical shift changes of peaks in ^15^N-^1^H HSQC experiments for each cocktail sample compared to a control containing only protein and 2.5% d_6_-DMSO (apo). Identification of compounds binding to the protein was achieved by screening the individual components from cocktails that gave rise to chemical shift changes.

#### Protein-ligand titrations

NMR experiments were conducted on a Bruker Avance 800 MHz spectrometer equipped with ^1^H, ^13^C, ^15^N triple resonance, Z-axis pulsed field gradient cryoprobe. ^15^N-^1^H HSQC spectra of 100 μM ^15^N NPAS2 PAS-A or 200 μM ^15^N CLOCK PAS-A WT/mutants were acquired in 180 μL volume in 3 mm NMR tubes with increasing concentration of KG- compounds (0, 100, 250, 350, 500, 750 μM) in NMR buffer (20 mM Tris-HCl pH 7.0, 50 mM NaCl, 2 mM TCEP), 2.5% d_6_-DMSO, and 10% D_2_O. All NMR data were collected at 25°C. NMR data were processed using NMRPipe/NMRDraw [66] and analyzed with NMRViewJ (One Moon Scientific) [67]. Chemical Shift Perturbations (Δδ) and binding constants (K_d_) were calculated using built-in titration analysis in NMRViewJ that utilized the following equations (1) and (2), respectively.

1. Δδ_TOT_ = [χ(Δδ^1^H)^2^ + (Δδ^15^N)^2^]^½^ and normalized with the scaling factor χ = 10
2. K_d_ was determined by fitting Δδ as a function of ligand concentration to the equation Y=(d((200+1/k+X)-((200+1/k+X)^2^-4[P]X) ^½^))/2[P], where Y = Δδ, d= maximum Δδ, X= ligand concentration, [P]= protein concentration, and 1/k = K_d_.

GraphPad Prism was used to plot binding curves [68].

#### WaterLOGSY NMR

NMR experiments were conducted on a Bruker Avance 800 MHz spectrometer, as above. ^1^H spectra of 500 μM KG-296 with 10 μM CLOCK or NPAS2 PAS-A were acquired in 180 μL volume in 3 mm NMR tubes in NMR buffer (20 mM Tris-HCl pH 7.0, 50 mM NaCl, 2 mM TCEP), 2.5% DMSO, and 10% D_2_O utilizing the pulse sequence *ephosgy* [44, 45]. Spectra were visualized in MestReNova (v15.1).

#### Protein backbone assignments

Backbone chemical shift assignments for NPAS2 PAS-A were obtained from BMRB entry 5019 [39]. The backbone assignment of mouse CLOCK PAS-A WT and the L170F mutant was accomplished using Bruker standard suite of triple resonance experiments HN(CA)CO, HNCO, HNCACB, CBCA(CO)NH, and C(CO)NH with modifications to permit non-uniform sampling (NUS) at 25% sampling. NMR experiments were acquired on Bruker Avance 800 MHz spectrometer equipped with ^1^H, ^13^C, ^15^N triple resonance, Z-axis pulsed field gradient cryoprobe. The NMR sample for assignment contained 600 µM ^13^C, ^15^N-labeled protein in a 330 μL sample volume in NMR buffer (20 mM Tris-HCl pH 7.0, 50 mM NaCl, 2 mM TCEP) in a Shigemi tube. POKY software was used for assignments [69].

#### High pressure NMR

High pressure NMR data were acquired on a Bruker Avance III HD NMR spectrometer at 700 MHz equipped with a 5 mm inverse TCI cryoprobe with pulsed-field Z-axis gradients. Samples were transferred to a zirconia tube of 3/5 mm inner/outer diameter connected to a syringe pump (Xtreme 60 pump, Daedalus Innovations LLC). A layer of mineral oil (270 μL) was added on top of the sample to separate it from the pressurizing liquid. Pressure NMR experiments were conducted with 300 μM protein (apo) or 300 μM protein + 1200 μM KG-296 (ligand-bound) in a baroresistant buffer containing a pair of buffer compounds (14.7 mM Tris pH 7.4, 35 mM sodium phosphate pH 7.4, 20 mM NaCl, and 20% D_2_O) to limit pressure-induced pH changes [70]. Sensitivity-enhanced ^15^N-^1^H HSQC spectra were acquired with pressure steps between 20-2500 bar, at increments of 250 bar, interleaving each high-pressure spectrum with a low pressure (20 bar) spectrum to establish the reversibility of changes in peak locations and intensities, allowing 10 min between pressure changes before acquiring data. Acquisition of each individual spectrum at a single pressure (high-pressure or interleaved 20 bar pressure) point took approximately 2 hours.

Peak intensities at every interleaved 20 bar HSQC after individual high pressure titration point were extracted for well-folded peaks/visible residues. Percent recovery was measured by taking the ratio of peak intensities at the beginning of pressure titration and after the end of titration (I_20bαr(before 250bαr point)_/I_20bαr(αfter 2250bαr point)_). The mean and standard deviation of the intensities were measured and plotted against pressure and time to generate a plot to compare protein stability over the whole titration series. One way ANOVA was run on the percent recovery data after the completion of the 2250 bar point using Prism (GraphPad). Statistical parameters, including sample size, precision measures (standard deviation, s.d.), and statistical significance are reported in Fig. 4 and corresponding figure legend.

### Protein crystallization

Human CLOCK PAS-A L170F was concentrated to 10 mg/ml and crystals were grown using the sitting drop vapor diffusion method in a 96-well plate using the Crystal Gryphon robotic nano-dispenser (ARI). Crystals grew in 2 μL of protein solution and 2 μL of reservoir solution (0.1 M MES pH 6.5, 1 M ammonium sulfate) and reached their maximum size within a week at room temperature. For apo CLOCK PAS-A WT/L170F, crystals were looped and soaked in a cryoprotectant solution (80% reservoir solution, 20% glycerol). For CLOCK PAS-A WT bound to KG-296, crystals were soaked in a cryoprotectant solution containing 1.5 mM KG-296 for 2 hours. All crystals were then flash-frozen in liquid nitrogen prior to data collection. Data sets were collected at the 5.0.3 beamline at the Advanced Light Source (ALS) at Lawrence Berkeley National Laboratory. Data were indexed, integrated, and merged using the CCP4 software suite [71]. Structures were determined by molecular replacement with Phaser MR82 using the apo structure of human CLOCK PAS-A (PDB: 6QPJ) [72]. Model building was performed with Coot [73], and structure refinement was performed with PHENIX [74]. All structural models and alignments were generated using PyMOL Molecular Graphics System 2.0 [75] and UCSF ChimeraX Molecular Visualization Program [76]. X-ray crystallography data collection and refinement statistics are provided in supplementary information (Table S1).

### Electromobility shift assay (EMSA)

*CLOCK:BMAL bHLH-PAS-AB binding to Cy5-E-box DNA* CLOCK-BMAL1 bHLH-PAS-AB WT or L170F (0-1000 nM protein) was titrated into a constant concentration of Cy5-labelled mouse *Per2* E1-box DNA (4 nM) (5’- [Cy5]GCGCGGTCACGTTTTCCAC-3’), synthesized and labeled by Sigma Aldrich. Reactions were performed in EMSA buffer (20 mM Tris-HCl pH 7.5, 75 mM NaCl, 10 mM KCl, 1 mM MgCl_2_, 0.1 mg/ml BSA and 1 mM DTT) and incubated at 4°C for 1 hour. After incubation, the samples were run by electrophoresis on a 4% native polyacrylamide gel in 0.5X TBE buffer (44.5 mM Tris, 44.5 mM Boric acid, and 1 mM EDTA). The bands were visualized by illuminating at 633 nm with a Typhoon TRIO imager (Amersham Biosciences).

#### Ligand competition assay

For KG-296 and KG-262 competition assays, CLOCK-BMAL1 bHLH-PAS-AB WT or L170F (100 nM) was incubated with Cy5-labelled E-box DNA (4 nM) as described above for 1 hour at 4°C. After which, increasing concentrations of the ligands (KG-296: 0-1 mM, KG-262: 0-1 mM) were added to the mixture and incubated at room temperature for 30 minutes. Samples were electrophoresed and imaged as described above. ImageJ was used for densitometry analysis, and model fitting was done using GraphPad Prism.

### Cell culture and assays

#### Cell culture

HEK293T cells were purchased from ATCC (cat. #CRL-3216) and cultured in 10% DMEM (i.e. 10% FBS) and 1X penicillin-streptomycin (Thermo Fisher) at 37°C in an incubator humidified with 5% CO_2_.

#### Per1-luciferase reporter assay

HEK293T cells were plated in a 48-well plate and transfected with LT-1 (Mirus) after 16- 20 hours with the following plasmids in octuple: for *Per1*-luc activity only: 4 ng pGL3 *Per1*- luciferase and 200 ng empty pcDNA4 vector; for CLOCK:BMAL1 activity: 4 ng pGL3 *Per1*- luciferase, 100 ng pSG5 *Clock* WT or L170F and 100 ng pSG5 *Bmal1*. After 24-30 hours, cells were harvested, lysed, and luciferase activity was measured with One-Glo EX Luciferase Assay System (Promega) on an EnVision plate reader (Perkin Elmer). Each reporter assay was repeated at least three times. Statistical parameters, including sample size, precision measures (standard deviation, s.d.), and statistical significance, are reported in Fig. 5 and the corresponding legend.

#### Western blotting

HEK293T cells were cultured as above and plated in 6-well plates. Cells were transfected with LT-1 (Mirus) in a 3 μL:1 μg LT-1:DNA ratio with 1 μg of pSG5 *Clock* WT or L170F and 1 μg pSG5 *Bmal1*. After 72 hours, cells were harvested, washed with PBS, and lysed in 200 μL of RIPA buffer (10 mM Tris-HCl, pH 8.0, 150 mM NaCl, 1 mM EDTA, 0.1% SDS, 0.1% Sodium Deoxycholate, 0.1% TritonX) containing complete protease inhibitor (Pierce) and phosphatase inhibitor tablets (Sigma). Following 15 minutes centrifugation at 19000 RPM at 4°C, the supernatant was added to 6x Laemmli buffer. A total of 15 μL of extracts/well were run on AnyKDa SDS-PAGE gels (Biorad) and transferred onto a nitrocellulose membrane and blocked for 1 hour at room temperature in TBST with 1% FBS. Western blotting was done using the following primary antibodies: the DYKDDDDK Tag antibody, HRP-conjugated (GenScript cat. # A01428-100) and β-actin Direct-Blot HRP antibody (BioLegend cat. # 664804). SuperSigna West Femto Maximum Sensitivity HRP substrate (Thermo Scientific cat. # 34095) was used for chemiluminescent detection on an imager (Biorad).

