## Supplemental information for "Small Molecule Regulation of CLOCK:BMAL1 DNA Binding Activity"

**Classification:** Biological Sciences; Biochemistry

**Keywords:** small-molecule screening, circadian rhythms, protein-ligand interactions, DNA binding, NMR spectroscopy, transcription factor

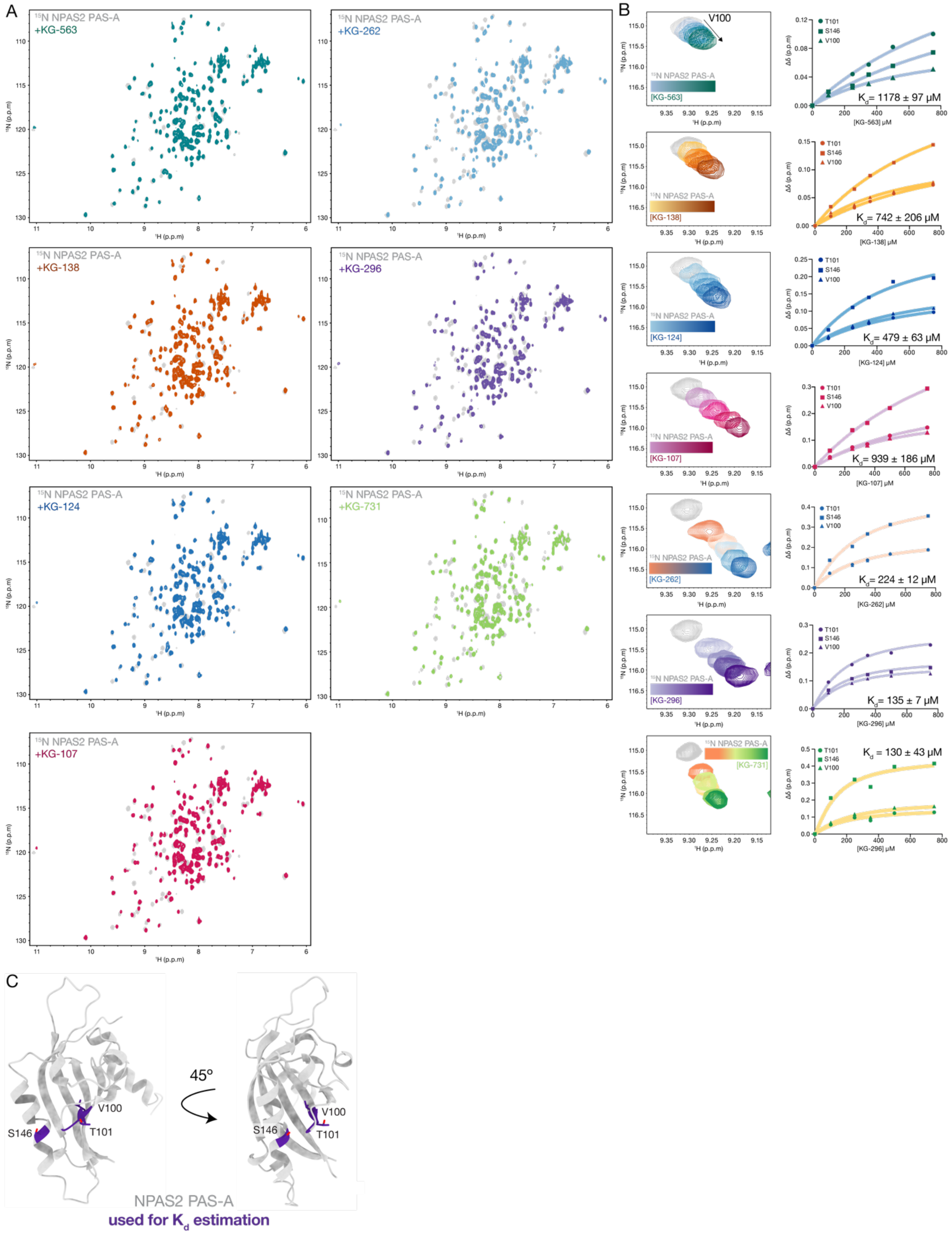


**Figure S1: NMR ligand screen of KG library identified 8 positive hits for NPAS2 PAS-A. (A)** $^{15}$N-$^{1}$H HSQC of $^{15}$N labeled NPAS2 PAS-A (gray) with 500 μM of KG ligands: KG-563 (dark green), KG-138 (brown), KG-124 (dark blue), KG-107 (pink), KG-262 (blue), KG-296 (purple), KG-731 (green). **(B)** Zoomed view of $^{15}$N-$^{1}$H HSQC of NPAS2 PAS-A V100 (apo, gray) showing dose dependent response of KG ligands (100-750 µM) in $^{15}$N labeled NPAS2 PAS-A, KG-563 (light blue to dark green), KG-138 (yellow to brown), KG-124 (light blue to dark blue), KG-107 (light pink to dark pink), KG-262 (peach to dark blue), KG-296 (purple to dark purple), KG-731 (orange to green). **(C)** NPAS2 PAS-A (gray, AlphaFold) showing 3 residues in sticks (purple) used for CSP fitting and apparent $K_d$ estimation.

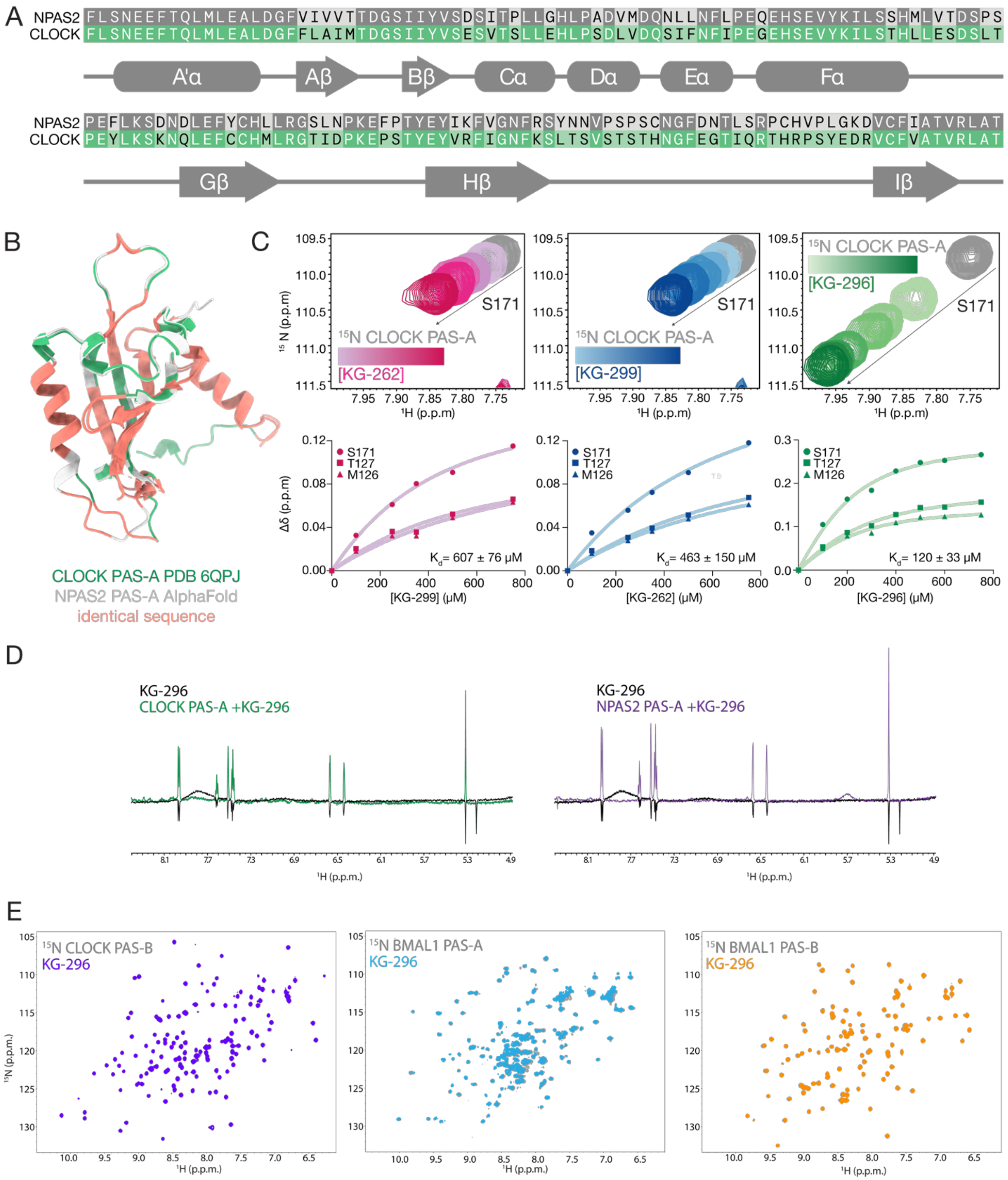
A
NPAS2 FLSNEEFTQLMLEALDGFVIVVTTDGSIIYVSDSITPLLGHLPADVMDQNLLNFLPEQEHSEVYKILSSHMLVTDSPS
CLOCK FLSNEEFTQLMLEALDGFFLAIMTDGSIIYVSESVTSLLEHLPSDLVDQSIFNFIPEGEHSEVYKILSTHLLESDSLT
A'α Aβ Bβ Cα Dα Eα Fα
NPAS2 PEFLKSDNDLEFYCHLLRGSLNPKEFPTYEYIKFVGNFRSYNNVPSPSCNGFDNTLSRPCHVPLGKDVCFIATVRLAT
CLOCK PEYLKSKNQLEFCCHMLRGTIDPKEPSTYEYVRFIGNFKSLTSVSTSTHNGFEGTIQRTHRPSYEDRVCFVATVRLAT
Gβ Hβ Iβ
B
CLOCK PAS-A PDB 6QPJ
NPAS2 PAS-A AlphaFold
identical sequence
C
15N CLOCK PAS-A
[KG-262]
S171
15N CLOCK PAS-A
[KG-299]
S171
15N CLOCK PAS-A
[KG-296]
S171
15N (p.p.m)
1H (p.p.m)
S171
T127
M126
Δδ (p.p.m)
Kd= 607 ± 76 μM
[KG-299] (μM)
Kd= 463 ± 150 μM
[KG-262] (μM)
Kd= 120 ± 33 μM
[KG-296] (μM)
D
KG-296
CLOCK PAS-A +KG-296
KG-296
NPAS2 PAS-A +KG-296
1H (p.p.m.)
E
15N CLOCK PAS-B
KG-296
15N BMAL1 PAS-A
KG-296
15N BMAL1 PAS-B
KG-296
15N (p.p.m.)
1H (p.p.m.)

F

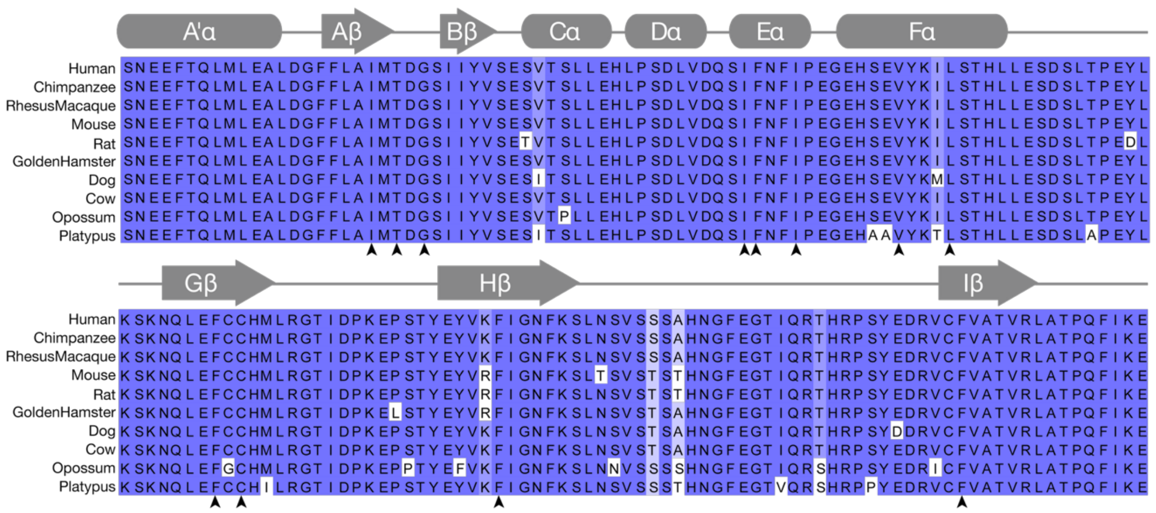
A'α
Aβ
Bβ
Cα
Dα
Eα
Fα
Human
Chimpanzee
RhesusMacaque
Mouse
Rat
GoldenHamster
Dog
Cow
Opossum
Platypus
Gβ
Hβ
Iβ

G

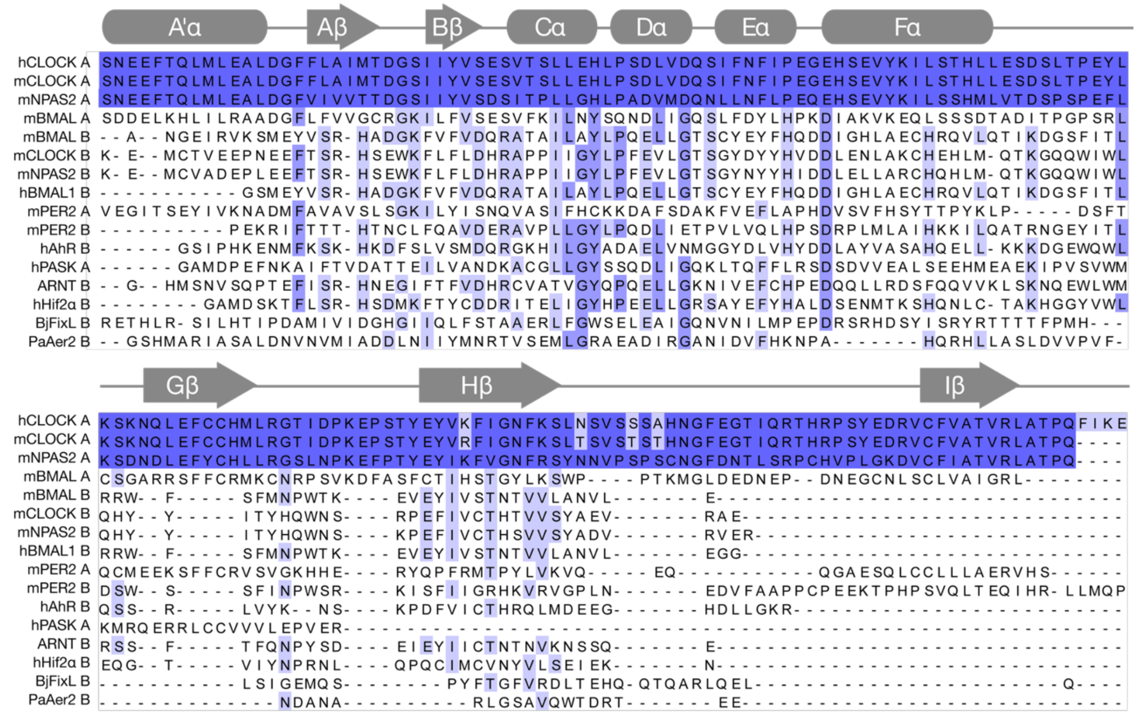
A'α
Aβ
Bβ
Cα
Dα
Eα
Fα
hCLOCK A
mCLOCK A
mNPAS2 A
mBMAL A
mBMAL B
mCLOCK B
mNPAS2 B
hBMAL1 B
mPER2 A
mPER2 B
hAhR B
hPASK A
ARNT B
hHif2α B
BjFixL B
PaAer2 B
Gβ
Hβ
Iβ

H

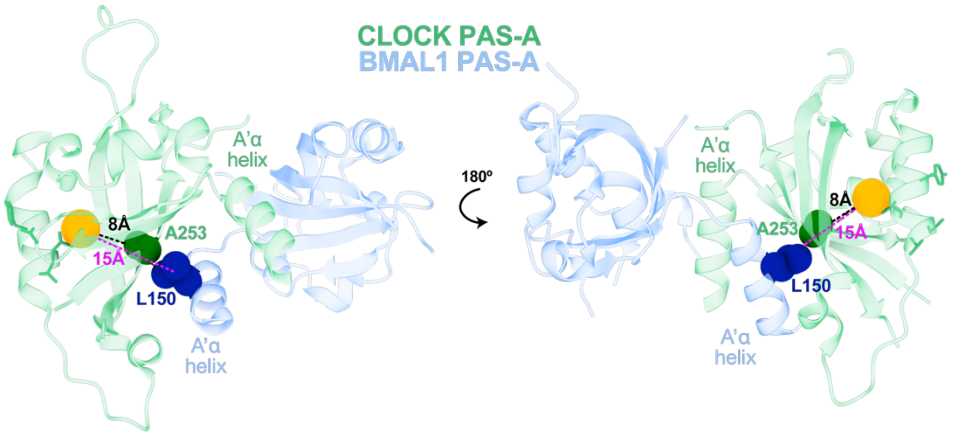
CLOCK PAS-A
BMAL1 PAS-A
A'α helix
8Å
A253
15Å
L150
A'α helix
180°
A'α helix
8Å
A253
15Å
L150
A'α helix

**Figure S2: CLOCK PAS-A and NPAS2 PAS-A bind KG compounds similarly. (A)** Sequence alignment of human NPAS2 PAS-A (gray) and CLOCK PAS-A (green). Darker shades indicate identical residues. Secondary structures of NPAS2 and CLOCK PAS-A (above). **(B)** Structure overlay of CLOCK (green, PDB 6QPJ) and NPAS2 PAS-A (gr*ay*, AlphaFold). Identical residues are shown in salmon. **(C)** Zoomed view of $^{15}$N-$^{1}$H HSQC of S171 in CLOCK PAS-A (apo, gray) showing dose-dependent response (100-750 μM) of KG-262 (pink), KG-299 (blue) and KG-296 (purple). Fitting concentration-dependent CSPs on $^{15}$N labeled CLOCK PAS-A to estimate apparent $K_d$ (bottom). **(D).** $^{1}$H WaterLOGSY spectra of 500 μM KG-296 (gray) with 10 μM CLOCK PAS-A (green, left) and 10 μM NPAS2 PAS-A (purple, right). **(E)** $^{15}$N-$^{1}$H HSQC spectra of $^{15}$N labeled CLOCK PAS-B (gray) with 500 μM KG-296 (violet), $^{15}$N BMAL1 PAS-A (gray) with 500 μM KG-296 (sky blue), $^{15}$N labeled BMAL1 PAS-B (gray) with 500 μM KG-296 (orange). **(F)** Sequence alignment of the CLOCK PAS-A domain from different species. Arrowheads represent residues showing maximum CSPs in Fig. 2D. **(G)** Alignment of PAS domains from proteins with reported endogenous or exogenous small-molecule ligands. 'A' and 'B' following the protein name indicate PAS-A and PAS-B, respectively. (A,B) Percent identity is shown by dark to light blue color for high to low conservation. CLOCK PAS-A was used as reference sequence for alignment in Jalview. **(H)** Distance measurements between the CLOCK PAS-A cavity center and the dimerization interface with BMAL1 PAS-A.

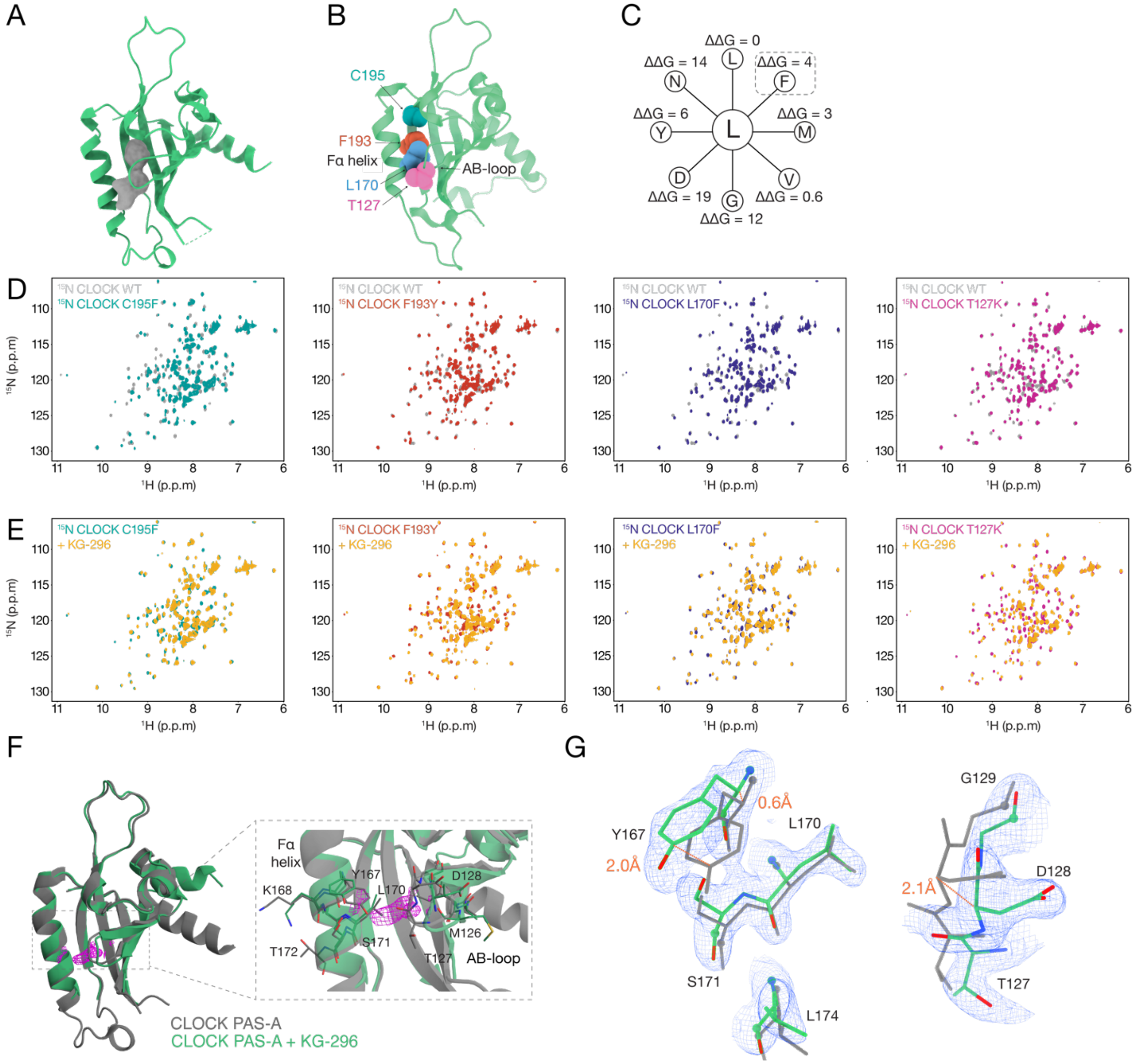


**Figure S3: Screening mutants to disrupt KG-296 binding to CLOCK PAS-A. (A)** CLOCK PAS-A (PDB 6QPJ, green) illustrating computationally mapped buried cavity (gray). **(B)** CLOCK PAS-A structure (PDB 6QPJ, green) highlighting location of mutants screened for KG-296 binding. **(C)** Pythia schematic showing some examples of L170 mutants and their ΔΔG scores in kcal/mol. **(D)** Overlay of $^{15}$N-$^{1}$H HSQC of $^{15}$N CLOCK PAS-A WT (gray) with $^{15}$N labeled CLOCK PAS-A mutants, C195F, teal; F193Y, brown; L170F, dark blue; T127K, pink. **(E)** $^{15}$N-$^{1}$H HSQC spectra of $^{15}$N CLOCK PAS-A mutants with 500 μM KG-296, as colored in (D) for apo, yellow with ligand. **(F)** Structure overlay of apo CLOCK PAS-A (gray, PDB:6QPJ) with CLOCK PAS-A + KG296 (green) showing experimental density near KG-296 from a $2mF_o\text{-}DF_c$ simulated-annealing composite omit map contoured at 1σ. **(G)** Zoom view of residues with change in backbone conformation with KG-296 incubation (green) relative to apo (gray).

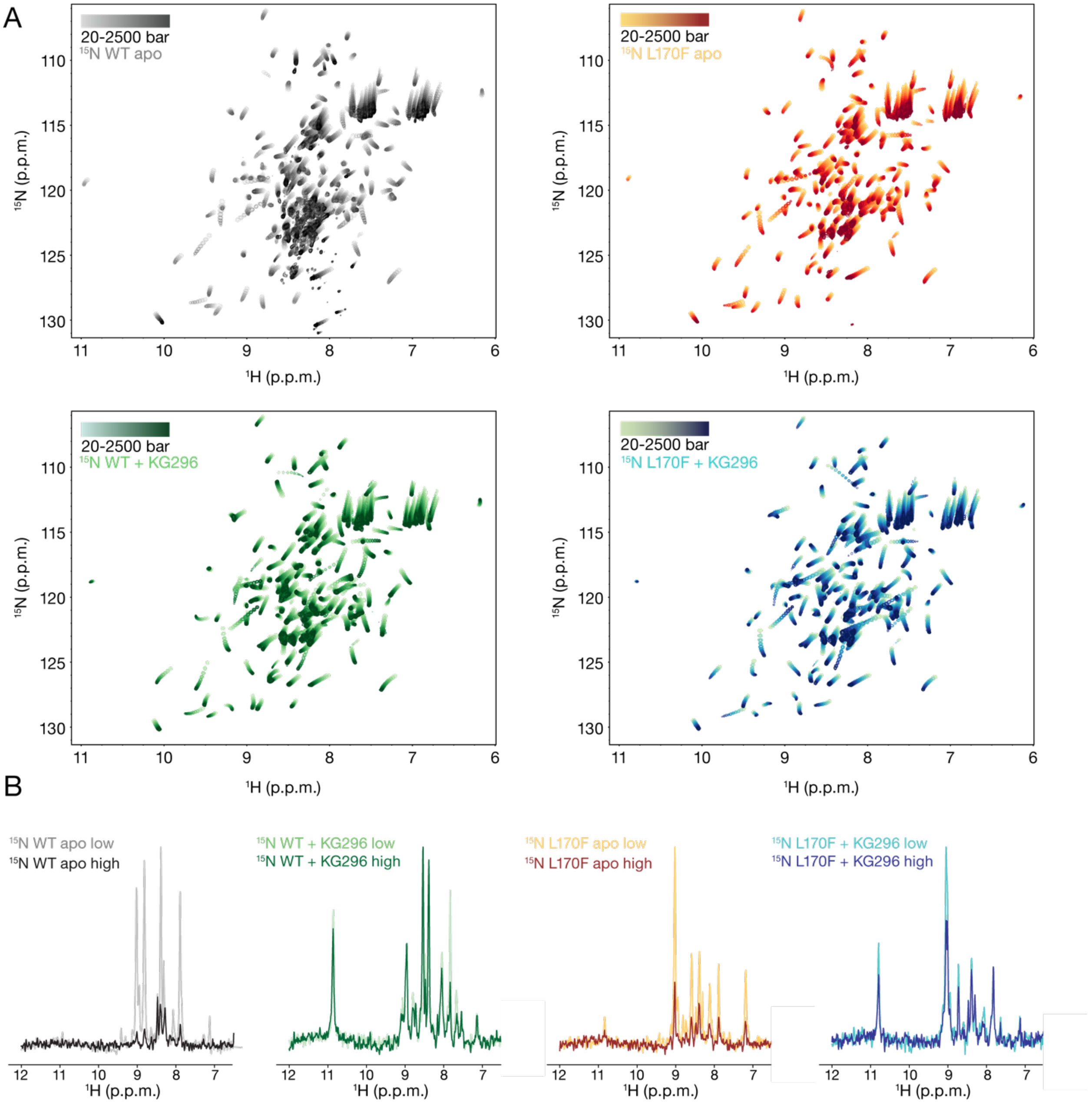


**Figure S4: High-pressure NMR titration reveals changes in CLOCK PAS-A due to KG-296 and/or L170F mutation. (A)** Overlays of $^{15}$N-$^{1}$H HSQC spectra acquired under increasing pressure from 20-2500 bar of CLOCK PAS-A apo (gray to black, top left), CLOCK PAS-A + KG296 (light green to dark green, bottom left), CLOCK PAS-A L170F apo (yellow to maroon, right left), CLOCK PAS-A L170 + KG-296 (light turquoise to dark blue, bottom right). (B). Overlay of $^{1}$H 1D traces extracted from $^{15}$N-$^{1}$H HSQC spectra at $^{15}$N = 118 ppm at 20 bar beginning of pressure titration point 250 bar (low) and at 20 bar after end of pressure titration point 2250 bar (high). WT apo (low, gray; high, black), CLOCK PAS-A + 1200 μM KG-296 (low, light green; high, dark green), L170F apo (low, yellow; high, maroon), L170F + 1200 μM KG-296 (low, turquoise; high, dark blue).

**Table S1.** X-ray crystallography data collection and refinement statistics

| Data collection | hCLOCK PAS-A L170F |
|---|---|
| PDB ID | 9PTN |
| Resolution range | 38.92 - 1.75 (1.813 - 1.75) |
| Space group | P 1 21 1 |
| Unit cell | 45.85 45.2101 76.63 90 92.97 90 |
| Total reflections | 307651 (30689) |
| Unique reflections | 31391 (3077) |
| Multiplicity | 9.8 (9.9) |
| Completeness (%) | 97.11 (97.31) |
| Mean I/sigma(I) | 40.73 (5.08) |
| R-merge | 0.5484 (0.7164) |
| CC1/2 | 0.792 (0.826) |
| Refinement statistics | |
| R-work | 0.1987 |
| R-free | 0.2179 |
| Number of non-hydrogen atoms | 2335 |
| macromolecules | 2128 |
| solvent | 207 |
| RMS(bonds) | 0.007 |
| RMS(angles) | 1.03 |
| Ramachandran favored/allowed (%) | 100 |
| Ramachandran outliers (%) | 0.00 |
| Average B-factor | 23.59 |
| macromolecules | 22.89 |
| solvent | 30.71 |

Statistics for the highest-resolution shell are shown in parentheses.
